# Loss of TLE3 Promotes Mitochondrial Program in Beige Adipocytes and Improves Glucose Metabolism

**DOI:** 10.1101/409748

**Authors:** Stephanie Pearson, Anne Loft, Prashant Rahbhandari, Judith Simcox, Sanghoon Lee, Anthony Donato, Peter Tontonoz, Susanne Mandrup, Claudio J. Villanueva

## Abstract

Prolonged cold exposure stimulates the recruitment of beige adipocytes within white adipose tissue. Beige adipocytes depend on mitochondrial oxidative phosphorylation to drive thermogenesis. The transcriptional mechanisms that promote remodeling in adipose tissue are not well understood. Here we demonstrate that the transcriptional coregulator TLE3 is induced with aging and inhibits mitochondrial gene expression in beige adipocytes. Conditional deletion of TLE3 in adipocytes prevents age- and diet-induced weight gain by promoting mitochondrial oxidative metabolism and increasing energy expenditure, thereby improving glucose control. Using chromatin immunoprecipitation and deep sequencing we found that TLE3 occupies distal enhancers in proximity to nuclear-encoded mitochondrial genes and that many of these enhancers are also enriched for EBF transcription factors. TLE3 interacts with EBF2 and blocks its ability to promote the thermogenic transcriptional program. Collectively, these studies demonstrate that TLE3 mediates age-dependent beige adipose thermogenic decline through inhibition of EBF2 transcriptional activity. Inhibition of TLE3 may provide a novel therapeutic approach for obesity and diabetes.

## Introduction

Subcutaneous white adipose tissue undergoes substantial remodeling in response to nutrient availability and changes in ambient temperature. Cold exposure promotes the recruitment of thermogenic adipocytes–a long-term adaptive mechanism that facilitates thermogenesis (1). Two distinct subtypes of thermogenic adipocytes have been found in mammals: brown and beige (2, 3). Many similarities exist between the two subtypes, including the presence of small, multilocular lipid droplets, and the characteristic expression of the mitochondrial uncoupling protein 1 (UCP1) (4, 5). UCP1 transports protons across the inner mitochondrial membrane without utilizing this potential energy for ATP synthesis (6), causing the chemical energy of substrates consumed by the cell in the TCA cycle to be released as heat (7). Beige adipocytes are also capable of generating heat through a futile creatine kinase cycle (8). Interest in brown adipose tissue (BAT) was catalyzed in the early 2000s, when advancements in positron-emission tomography (PET) scanning demonstrated the presence of BAT in adult humans (9-12).

Despite their functional similarities, beige adipocytes arise from a separate lineage from brown adipocytes and are distinctively located as clusters of cells within white adipose tissue (WAT) depots (3, 13, 14). Not all white adipose tissue is prone to acquire beige adipocytes: subcutaneous depots such as the inguinal white adipose tissue (iWAT) recruit beige adipocytes in response to long-term cold exposure or after stimulation β3-adrenergic receptor agonists (15), while much of the visceral adipose tissue, such as the epididymal WAT (eWAT) is resistant to beiging. There is also evidence that as mice age, the conversion of beige adipocytes within white adipose tissue becomes impaired, although the underlying mechanisms are unknown (16).

There has been considerable dispute over the identity of thermogenic tissue in adult humans, but mounting evidence points to the existence of BAT (17), as well as populations of cells bearing markers of beige adipocytes (3, 18). Pathological conditions such as pheochromocytoma that lead to excess catecholamine production are associated with a greater abundance of beige adipocytes that is thought to promote a hypermetabolic state (19, 20). Since the presence of these thermogenic tissues are inversely related to metabolic disorders and obesity (7, 21), understanding the regulatory mechanisms that govern the development and activity of thermogenic adipocytes has become an area of intense therapeutic interest.

All adipocyte subtypes require peroxisome proliferator-activated receptor gamma (PPARγ), which is necessary and sufficient to drive adipocyte differentiation (22). A prior genome-wide survey of PPARγ binding profiles from primary cells isolated from three separate tissues (iWAT, eWAT, and BAT) differentiated *in vitro* showed an overlap in binding and similar intensities at the majority of PPARγ binding sites (23). However, this study also revealed a subset of depot-specific binding sites that correlated well with depot-specific gene expression (23). These findings suggested that PPARγ likely works in concert with other transcription factors and transcriptional co-regulators to regulate cell-type specific gene expression, although the identities of these auxiliary factors remain to be fully elucidated.

Among the transcription factors known to work in concert with PPARγ to determine adipocyte identity are those of the early B-cell factor (EBF) family. Both EBF1 and EBF2 participate in early adipogenesis (24, 25), and it has recently been demonstrated that they also play important roles in differentiated adipocytes. EBF1 positively regulates insulin-stimulated glucose uptake and lipogenesis (26), (27), while EBF2 is selectively expressed in brown adipocytes and facilitates the binding of PPARγ to brown fat-specific genes to promote thermogenesis (28)-(29). The EBF DNA binding motif associates with BAT-specific PPARγ binding sites (28) and enhances expression of genes important for mitochondrial function (30).

The transducin-like enhancer of split (TLE) protein family is a highly conserved group of transcriptional co-regulators (31), (32). Groucho, the TLE orthologue in *D. Melanogaster*, was first identified as a key regulator of neuronal development (31). A high-throughput screen identified TLE3 as a PPARγ cofactor that promotes white-specific gene expression (33, 34). TLE3 has been shown to disrupt the interaction between Prdm16 and PPARγ, causing the BAT of *Tle3* transgenic mice to appear more like white adipose tissue, with larger lipid droplets and higher levels of WAT-selective transcripts (34). During preadipocyte differentiation TLE3 activates PPARγ to promote differentiation of all adipocyte subtypes, and to sustain the lipid-storage transcriptional program after differentiation. TLE3 is expressed in all adipocytes, but its expression is highest in WAT when compared to BAT.

Here we report that conditional deletion of TLE3 in adipocytes promotes the recruitment of beige adipocytes, increases energy expenditure, and leads to improved glucose metabolism and insulin sensitivity. Analysis of genome-wide occupancy of TLE3 in adipocytes suggests a direct role for this cofactor in regulating the nuclear-encoded mitochondrial gene expression program. Moreover, TLE3 binds to genomic regions that also associate with EBF transcription factors. We define a functional interaction between TLE3 and EBF2 in which TLE3 attenuates the EBF2-driven transcriptional program. Our findings outline a transcriptional mechanism that governs mitochondrial gene expression and the age-dependent programing of beige adipocytes.

## Results

### Conditional Deletion of TLE3 in Adipocytes Promotes a Beige Transcriptional Program

To test the impact of TLE3 on whole body energy metabolism, we generated mice with a conditional deletion of TLE3 in adipocytes. We acclimated the *Tle3*^F/F^ Adiponectin-Cre (Adipo-TLE3 KO) mice and their littermate controls, aged 3-4 months, to one of two temperature conditions— thermoneutrality (30°C) or cold exposure (4°C)—for 7 days. At the conclusion of the study, the iWAT, eWAT, and BAT depots were fixed, sectioned, and stained with Hematoxylin and Eosin (H&E). Under thermoneutral conditions, we observed no differences in adipose histology between control and Adipo-TLE3 KO mice (**Figure 1A, top panel**). However, when the mice were subjected to cold exposure, both groups had reduced lipid droplet size in iWAT, but strikingly, there were more multilocular cells consistent with beige adipocytes in the Adipo-TLE3 KO mice (**Figure 1A, bottom panel**). Both eWAT and BAT showed similar histological features between control and Adipo-TLE3 KO mice in the setting of cold exposure (**Figure 1A**). No alterations were observed in liver morphology at thermoneutrality or at 4°C (**Figure S1A**). We measured gene expression within iWAT following 4-days of cold exposure and found increases in thermogenesis-associated genes, such as *Cited, Dio2, Elovl3, Eva1, Tmem26* and *Ucp1*, while the genes encoding general adipocyte markers such as adiponectin and aP2 remained unchanged (**Figure 1B**). These changes were specific to iWAT, as no changes in gene expression were observed between control and Adipo-TLE3 KO mice in the eWAT or BAT depots (**Figure S1B**).

**Figure 1:**
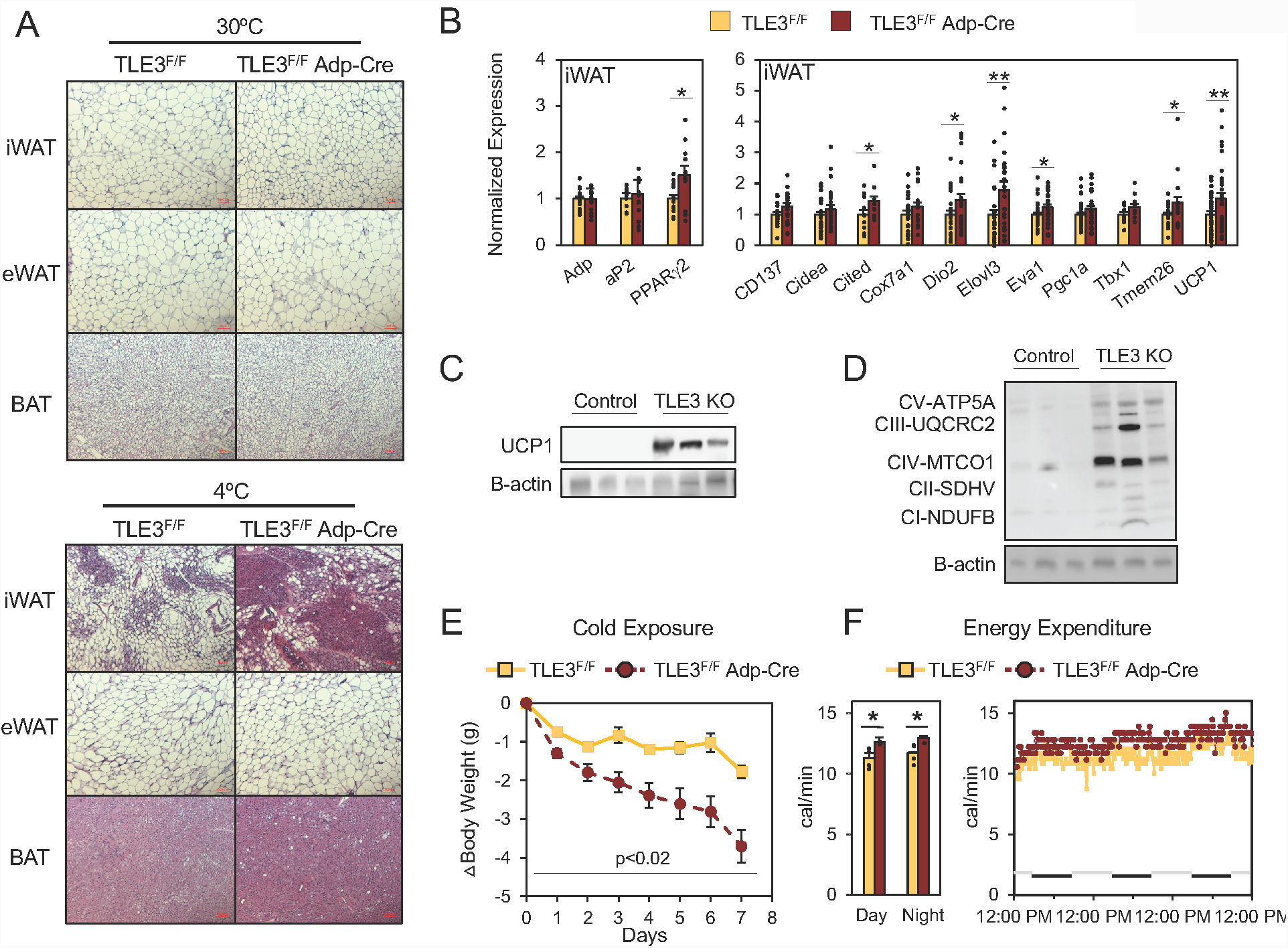
Conditional deletion of TLE3 in adipocytes promotes beige adipocyte programming and whole-body energy expenditure. **(A)** Histological analysis of iWAT, eWAT, and BAT at thermoneutrality (30°C) and cold (4°C) for control (TLE3^F/F^) and Adipo-TLE3 KO (TLE^F/F^ Adp-Cre) mice. Scale bar, 500 μm. **(B)** Expression of adipogenic and thermogenic-associated genes from the iWAT measured by real-time PCR after 4 days cold exposure. Normalized to *36B4*. **(C)** UCP1 protein expression in iWAT after 4 days cold exposure. **(D)** Protein expression for oxidative phosphorylation proteins in iWAT after 4 days cold exposure. **(E)** Time-dependent changes in body weight for control and Adipo-TLE3 KO mice housed at 4°C (n = 4/group). **(F)** Energy expenditure as measured by indirect calorimetry in the comprehensive lab animal monitoring system (CLAMS) metabolic cages for control and Adipo-TLE3 KO mice (n = 4/group). Data presented as mean ± SEM. Significance analyzed by two-tailed Student’s t-test, *p<0.05, and **p<0.01.

The greater recruitment of beige adipocytes was further verified by the finding of increased UCP1 protein in iWAT of Adipo-TLE3 KO mice (**Figure 1C**). As beige adipocytes are rich in mitochondria, we also measured mitochondrial content by blotting for proteins from each of the electron transport chain complexes. The iWAT from Adipo-TLE3 KO mice expressed more protein from each subunit, with marked increases in the cytochrome oxidase, complex IV (**Figure 1D**).

### Loss of TLE3 in Adipocytes Increases Whole-Body Energy Expenditure

We next proceeded to monitor the animal during cold exposure and we observed greater weight loss over time in the Adipo-TLE3 KO group, compared to the controls (**Figure 1E**). To assess whether these changes in body weight were due to decreased food consumption or increased energy expenditure, we placed age-matched mice in metabolic cages at 4°C for a three-day period. We observed no differences in body weight or body composition between the two groups prior to cold exposure (**Figure S1C**). Adipo-TLE3 KO mice also showed an increase in energy expenditure in both day and night when compared to the control group (**Figure 1F**). Measurements of O_2_ consumption and CO_2_ production (**Figure S1D**) revealed a modest but consistent increase in the cold. During this period we did not find measurable differences in food consumption or activity (**Figure S1E**).

### TLE3 Regulates Glucose Utilization and Insulin Sensitivity

We next assessed whether having more beige adipocytes in the setting of TLE3 deletion promoted glucose utilization. As increased beige and brown adipose tissue have previously been linked to improvements in glucose handling (35, 36) we asked whether the Adipo-TLE3 KO mice had improved glucose clearance. We performed glucose tolerance tests (GTTs) on Adipo-TLE3 KO and control mice that had been acclimated to thermoneutrality for one week, followed by 4 days of cold exposure and a second GTT. No differences were observed in the glucose excursion curves at 30°C. As expected, both groups demonstrated improvements after cold acclimation (**Figure 2A**), but the change was greatest in the Adipo-TLE3 KO group (**Figure 2B**). We performed an insulin tolerance test (ITT) following the same procedure, and while the change to the control group was modest, insulin sensitivity of Adipo-TLE3 KO mice was improved following cold exposure (**Figure 2C-D**). No differences were observed in plasma insulin levels between the two groups after cold exposure (**Figure S2A**). Because our metabolic phenotyping involved acclimation to 4°C (a temperature that triggers shivering), we recognized that glucose clearance into muscle could contribute to these effects. Thus, to control for shivering-induced glucose clearance, we performed a dosing regimen for both control and Adipo-TLE3 KO groups with a β3-adrenergic receptor agonist, CL-316,243, at 1 mg/kg/day for 3 days to stimulate the thermogenic gene program in adipose tissue. We found that following CL-316,243 administration, the fasting blood glucose levels had greater improvements in the Adipo-TLE3 KO mice than the control group (**Figure 2E**).

**Figure 2:**
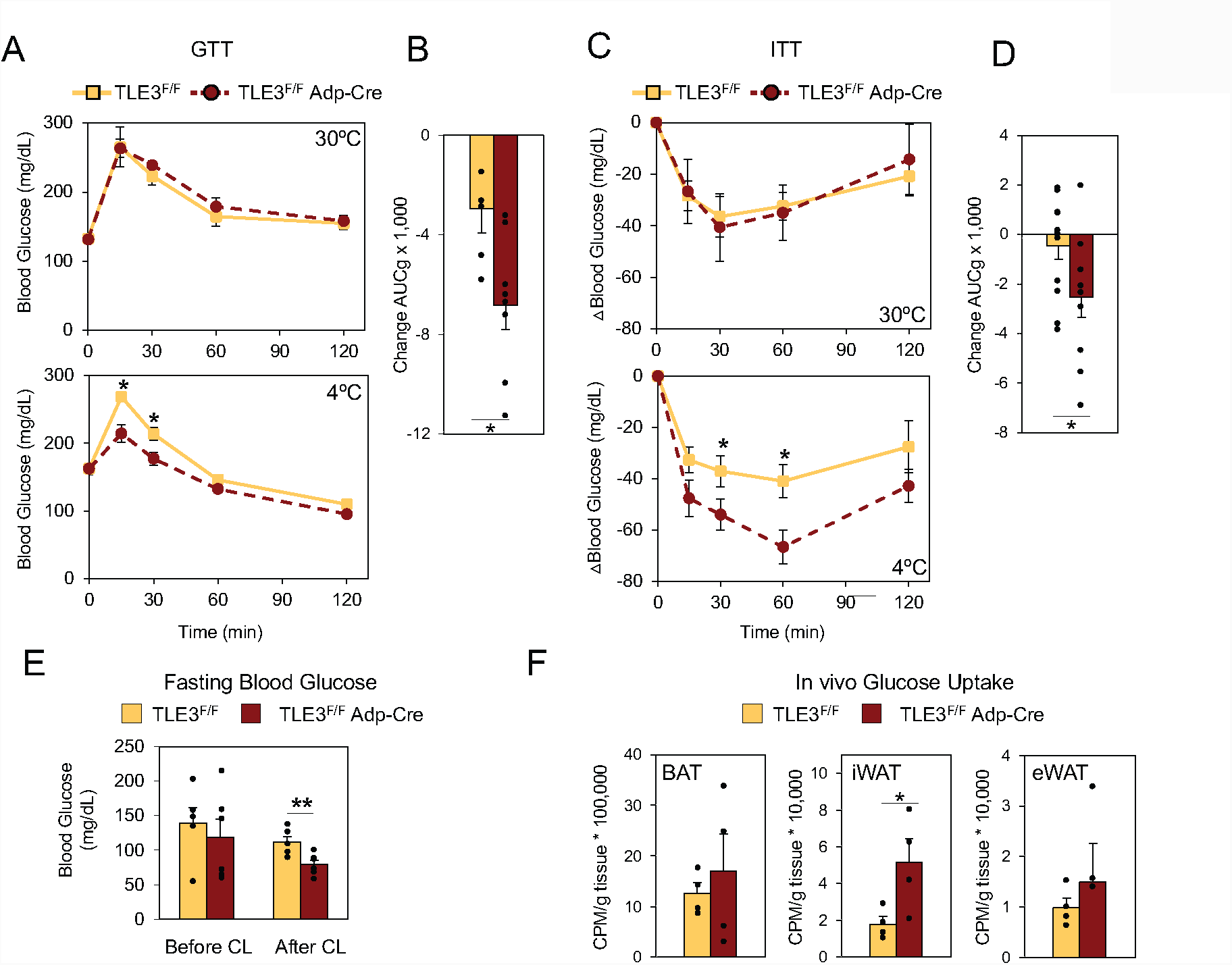
Deletion of TLE3 in adipocytes improves glucose handling with cold exposure. **(A).**Glucose tolerance test (GTT) on control (TLE3^F/F^) and Adipo-TLE3 KO (TLE3^F/F^ Adp-Cre) mice after 1 week at thermoneutrality (30°C) and 4 days in cold (4°C) (n = 6-7/group). **(B)** Change in measured area under the curve for the blood glucose levels (AUC_g_) from GTT performed after thermoneutrality or after cold exposure (n = 6-7/group). **(C)** Insulin tolerance test (ITT) on control and Adipo-TLE3 KO mice after 1 week at thermoneutrality and cold (n = 6-7/group). **(D)** Change in measured area under the curve for the blood glucose levels (AUC_g_) from ITT performed after thermoneutrality to GTT performed after cold exposure (n = 6-7/group). **(E)** Fasting blood glucose levels of control and Adipo-TLE3 KO mice before and after administration of 1 mg/kg CL-316,243 daily for 3 days (n = 5-6/group). **(F)** Uptake of 2-[U-^14^C]-deoxyglucose in BAT, iWAT, and eWAT for control and Adipo-TLE3 KO mice following 4 days of cold exposure. Data presented as mean ± SEM. Significance analyzed by two-tailed Student’s t-test, *p<0.05, and **p<0.01.

Under these conditions, beige adipocytes represent a small fraction of the total adipocyte content of the body. This led us to ask whether beige adipocytes were abundant enough to account for these changes in glucose handling. We observed increases in the expression of Glut1 and Glut4 transporters in the iWAT depot of Adipo-TLE3 KO mice, but not in the BAT or eWAT (**Figure S2B**), but this only indicates the tissue is poised to augment glucose clearance. To directly test the contribution of beige adipocyte to the phenotype of Adipo-TLE3 KO mice, we performed *in vivo* glucose uptake assays with uniformly-labelled ^14^C-2-deoxyglucose in mice housed at 4°C for 4 days. Thirty minutes following radiolabeled glucose administration, the animals were euthanized, tissues were excised, and the radioactive content of each tissue measured. Glucose uptake in TLE3-deficient iWAT was increased three-fold compared to controls, while only modest, nonsignificant increases were found in BAT and eWAT (**Figure 2F**). These data suggest that the changes in whole-body glucose handling observed in the Adipo-TLE3 KO mice are primarily driven by uptake into beige adipose tissue, and that increased beige adipose tissue alone is capable of regulating system glucose metabolism.

### TLE3 Intrinsically Regulates the Thermogenesis and Glucose Clearance in Beige Adipocytes

Beige adipocytes rely on both intrinsic regulation and systemic signaling to control their recruitment. To ascertain whether the phenotypic changes we were observing *in vivo* were due to alterations in TLE3-dependent gene transcription, we utilized immortalized TLE3^F/F^ iWAT preadipocytes with a tamoxifen-inducible Cre (pMSCV2-CreERT2). Because TLE3 is known to be involved with early adipocyte differentiation, we chose to knockout TLE3 four days after the differentiation cocktail was administered (33). To ensure that the differentiation potential was not affected by knocking out TLE3 at this timepoint, we stained for lipids using both Oil Red O and BODIPY and found similar lipid accumulation in TLE3-knockout cells when compared to controls (**Figure 3A**). In this *in vitro* system, the loss of TLE3 led to increased UCP1 protein but no change in PPARγ protein levels (**Figure 3B**). Further, these protein changes were accompanied by increases in other thermogenesis-associated gene transcripts, such as *Cidea, Dio*2, *Ppargc1*a, *Cox7a1*, and *Ucp1*, while no changes were observed in transcript levels of adipocyte differentiation markers such as *aP2* (**Figure 3C**). Similar to our *in vivo* findings, we found increased expression of the *Glut4* glucose transporter in the TLE3-knockout cells (**Figure S3A**). There was, however, no observable change in the thermogenesis genes due to tamoxifen administration in the empty vector (pMSCV2) cells (**Figure S3B**).

**Figure 3:**
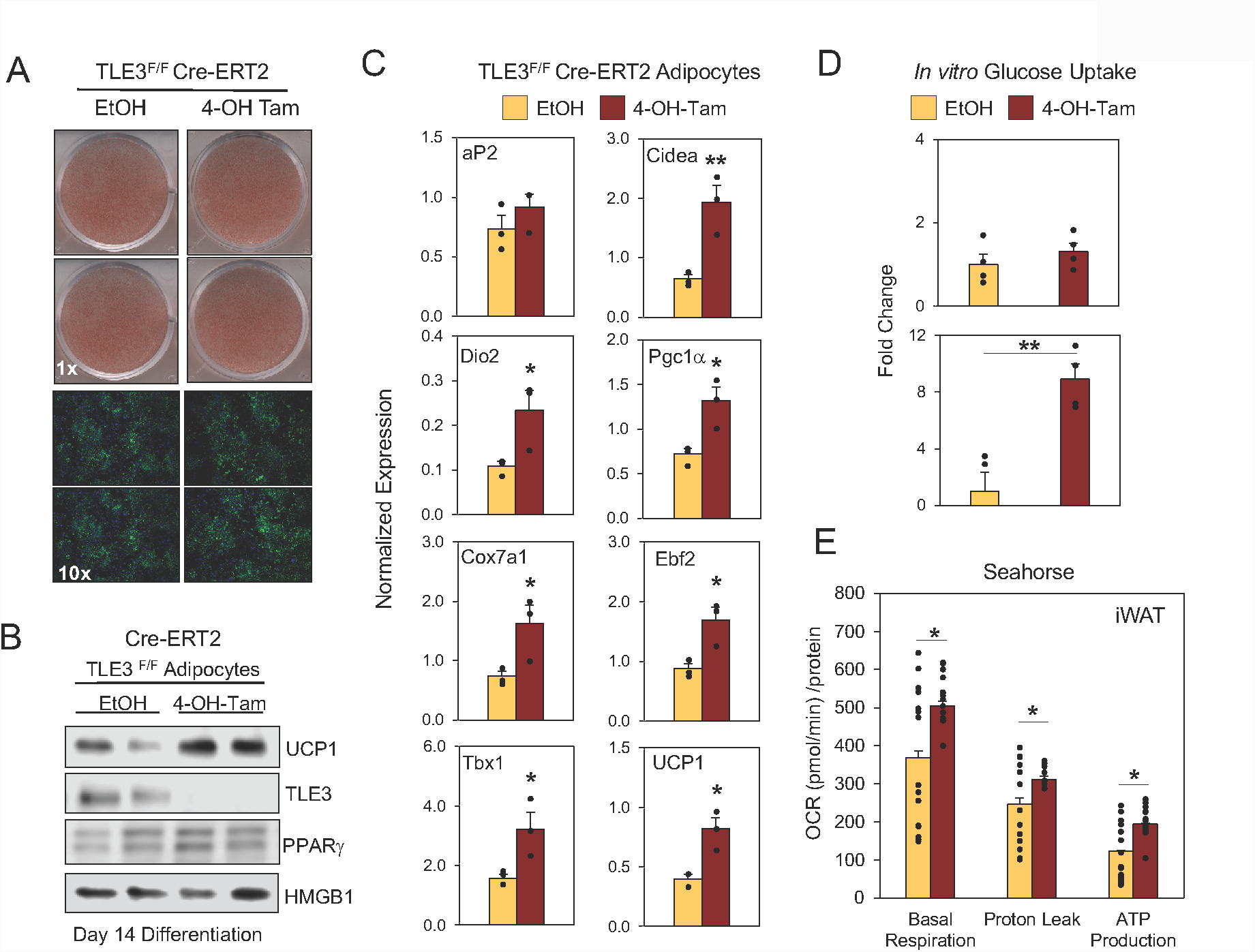
Deletion of TLE3 promotes the programming of beige adipocytes and glucose clearance *in vitro*. **(A)** Oil Red O staining (top) and DAPI/BODIPY staining (bottom) of TLE3^F/F^ iWAT preadipocytes expressing pMSCV2-CreERT2 and treated with either EtOH (control) or 4-hydroxytamoxifen on day 4 after administration of differentiation cocktail and stained on day 10. Protein expression of UCP1 and TLE3 by immunoblot in TLE3^F/F^ CreERT2 iWAT preadipocytes differentiated in culture. **(C)** Gene expression measured by real-time PCR in TLE3^F/F^ CreERT2 iWAT preadipocytes differentiated in culture. Data normalized to *36B4* (n = 3/group). **(D)** *In vitro* glucose uptake of 2-[U-^14^C]-deoxyglucose in TLE3^F/F^ iWAT preadipocytes, expressing either pMSCV2 (empty vector) or CreERT2 (n=4/group). **(E)** Oxygen consumption rate (OCR) of TLE3^F/F^ CreERT2 iWAT preadipocytes, as measured by Seahorse XF Analyzer (n=14/group). Data presented as mean ± SEM. Significance analyzed by two-tailed Student’s t-test, *p<0.05, and **p<0.01.

Having established an *in vitro* model for the beige adipocyte recruitment we observed *in vivo*, we next performed *in vitro* glucose uptake with uniformly-labelled ^14^C-2-deoxyglucose to determine whether the glucose clearance observed previously was due to intrinsic changes or whether TLE3-deficient iWAT was responding to systemic signaling. Upon the administration of tamoxifen to CreERT2-expressing cells, the loss of TLE3 causes approximately a 4-fold increase in glucose uptake, while no change was observed pMSCV2 cells treated with 4-OH-tamoxifen (**Figure 3D**). We also observed that the loss of TLE3 led to increases in oxygen consumption during basal respiratory conditions, isolated proton leak and ATP-production (**Figure 3E**), demonstrating a rise in energy utilization in beige adipocytes lacking TLE3.

### TLE3 Promotes Diet- and Age-induced Weight Gain and Glucose Intolerance

We appreciate that 4°C cold exposure represents a severe thermogenic stimulus, and we wanted to understand whether the loss of adipose TLE3 could affect systemic metabolism in the absence of a cold stimulus. Therefore, the Adipo-TLE3 KO mice and their littermate controls were placed on a high-fat diet (HFD, 60% fat; 5.2 kcal/g) for a period of 14 weeks and housed at ambient temperature (22-24°C). We monitored body weight weekly and found that the Adipo-TLE3 KO mice showed a degree of protection against weight gain (**Figure 4A**). Analysis of body composition revealed that the control group had higher body fat beginning after 6 weeks on HFD, and the difference was maintained until the end of the study (**Figure 4B**). To test if loss of TLE3 in this setting was also associated with improved glucose clearance, we performed a GTT. We observed elevated fasting glucose levels in both groups, but the Adipo-TLE3 KO mice showed enhanced glucose excursion (**Figure 4C**). An ITT showed similar insulin resistance between both groups, although the Adipo-TLE3 KO mice trended toward lower glucose levels (**Figure S4A**). Histology indicated that the Adipo-TLE3 KO mice also had more beige adipocytes than the control group on HFD (**Figure S4B**).

**Figure 4:**
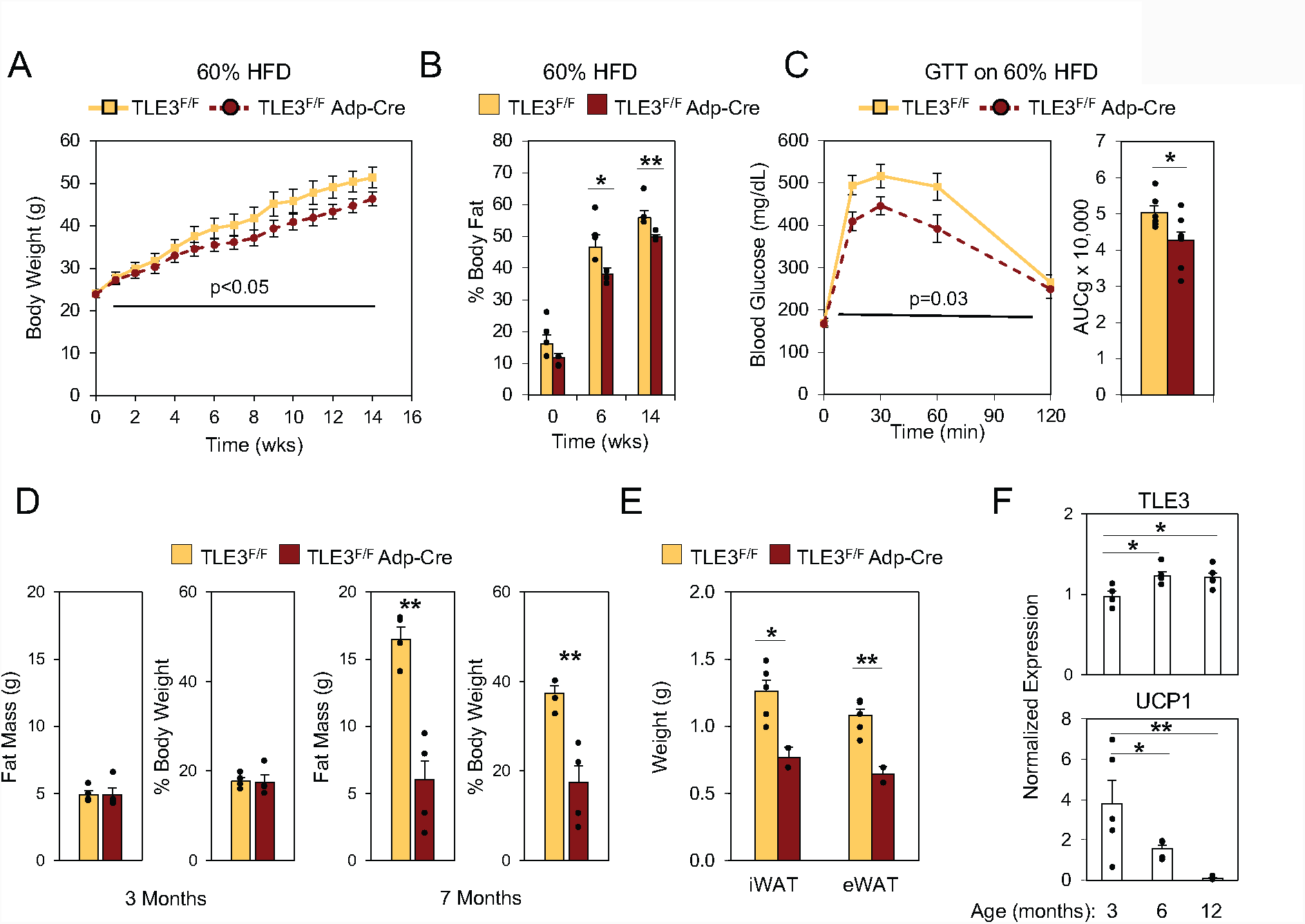
Conditional deletion of TLE3 in adipocytes protects against diet- and age-induced weight gain and glucose intolerance. **(A)** Time course of body weight for control (TLE3^F/F^) and Adipo-TLE3 KO (TLE3^F/F^ Adp-Cre) mice housed at ambient temperature on 60% high fat diet (HFD) for 14 weeks (n = 6-8/group). **(B)** Body fat composition by NMR in control and Adipo-TLE3 KO groups on 60% HFD over time (n = 6-8/group). **(C)** Glucose tolerance test (GTT) on control and Adipo-TLE3 KO mice after 14 weeks on 60% HFD (n = 6-7/group). **(D)** Body fat composition by nuclear magnetic resonance (NMR) in control and Adipo-TLE3 KO groups at 3 months and 7 months of age on chow diet, housed at ambient temperature (n = 6/group). **(E)** Weight of white fat depots excised from control and Adipo-TLE3 KO mice at 7 months of age on chow diet, housed at ambient temperature (n = 4-7/group). **(F)** Gene expression TLE3 and UCP1 in iWAT of wild type (C57BL/6) mice at 3, 6, and 12 months of age after 5 hours cold exposure (n = 5/group). Data presented as mean ± SEM. Significance analyzed by two-tailed Student’s t-test, *p<0.05, and **p<0.01.

We hypothesized that the improvements observed on HFD were due to the promotion of thermogenesis caused by the loss of TLE3. If true, this phenomenon should also be in play in Adipo-TLE3 KO mice on chow diet, but the phenotypic consequences might only become apparent after an extended period of time. Accordingly, we placed Adipo-TLE3 KO mice and their littermate controls on a normal chow diet (7% fat; 3.1 kcal/g), housed them at room temperature, and allowed them to age to 7 months. Consistent with our previous observations, at 3 months of age there were no observable differences in body weight, fat mass, or body fat percentage between the two groups (**Figure S1C**). However, when the control group experienced age-induced weight gain, the Adipo-TLE3 KO mice were protected (**Figure 4D**). Notably, at the time of sacrifice, both the iWAT and eWAT fat pads of the Adipo-TLE3 KO mice weighed significantly less than those of the control group (**Figure 4E**). Both eWAT and iWAT showed reduced adipose tissue size in Adipo-TLE3 KO mice when compared to controls (**Figure S4C**). We also observed increases in TLE3 expression in the iWAT of wild-type mice with age, which corresponded with a marked decrease in the expression of UCP1 (**Figure 4F**). Together these findings suggest TLE3 contributes to the impairment of age-associated thermogenesis and weight gain.

### Genome-wide Analysis Reveals Preferential Association of TLE3 with WAT Genes

TLE3 was initially identified as a transcriptional coactivator of PPARγ during early adipocyte differentiation (33). To develop a broader understanding of TLE3’s role in regulating gene expression, we performed genome-wide chromatin-immunoprecipitation in eWAT-, iWAT-, and BAT-derived adipocytes, followed by deep sequencing (ChIP-seq). We found that TLE3 had a greater number of bindings sites in WAT compared to BAT, both in terms of number of binding regions (**Figure 5A**) and binding intensity at each region (**Figure 5B**). These findings are consistent with the previous report of lower TLE3 expression in BAT (34). Although TLE3 mapped to many common sites within the three adipocyte subtypes, we also identified unique binding sites for each adipocyte subtype (**Figure 5A**).

**Figure 5:**
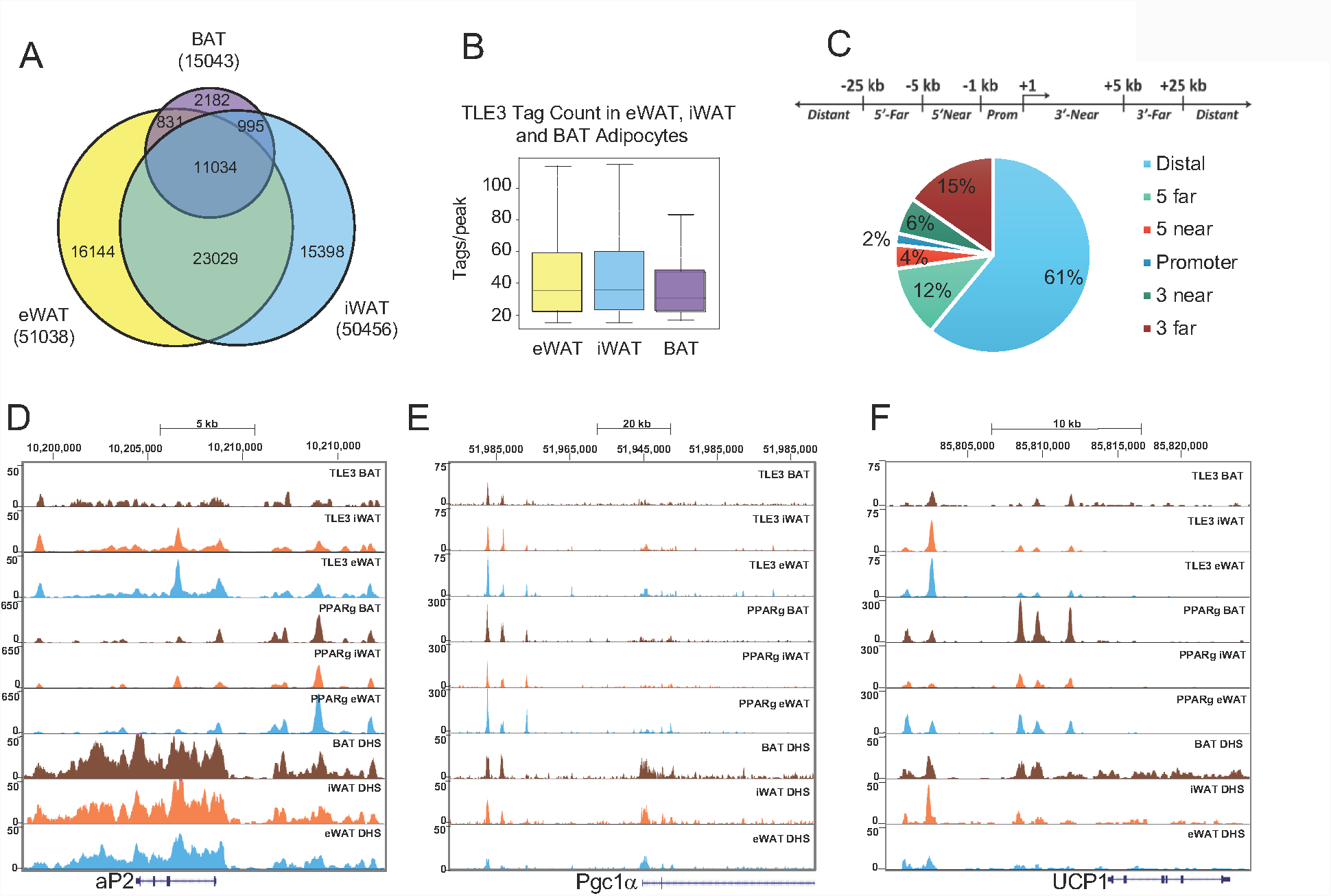
Genome-wide profiling of TLE3 in adipocytes reveals increased binding in WAT. **(A)** Venn diagram demonstrating number of TLE3 binding sites in eWAT, iWAT, and BAT. Filtered for >15 tags and >10-fold above input. **(B)** Binding intensity at sites occupied by TLE3 in eWAT, iWAT, and BAT. **(C)** Genomic distribution of TLE3 relative to the distance from transcriptional start sites (TSS). **(D)** Genome browser tracks of *aP2* for TLE3 and PPARγ in eWAT, iWAT, and BAT. **(E)** Genome browser tracks of *Pgc1α* for TLE3 and PPARγ in eWAT, iWAT, and BAT. **(F)** Genome browser tracks of *UCP1* for TLE3 and PPARγ in eWAT, iWAT, and BAT.

The genomic distribution of TLE3 mapped principally to distal regions of the mouse genome (**Figure 5C**), an observation that is consistent with the TLE/Groucho family being located primarily at enhancers (37, 38). There was nearly equal distribution of binding upstream and downstream of the transcriptional start site (TSS) (**Figure S5A**). Despite unique elements of TLE3 binding in each tissue, the distribution of TLE3 was largely similar between iWAT, eWAT, and BAT (**Figure S5B**). Through our genomic analysis we found that 61% of TLE3 binding sites are >25 kb distal from TSS. TLE3 was previously shown by ChIP-qPCR to occupy the aP2 enhancer/promoter. A closer examination of our ChIP-seq dataset revealed several TLE3 binding sites nearby the aP2 gene. Using data from a PPARγ ChIP-seq of differentiated progenitors of eWAT, iWAT, and BAT (23) we observed that TLE3 occupancy overlapped with multiple prominent PPARγ binding sites (**Figure 5D**). TLE3 was previously reported to interact with PPARγ at the promoters of key adipocyte target genes (33). To determine the overlap in binding sites between TLE3 and PPARγ we aligned TLE3 and PPARγ ChIP-seq datasets and found that 25% of TLE3 WAT-binding sites overlap with PPARγ, while 49% of PPARγ WAT-binding sites overlap with TLE3 (**Figure S5C**). PPARγ WAT binding sites were defined as PPARγ bindings sites shared between eWAT and iWAT. Examples of the distribution of TLE3 binding sites relative to PPARγ can be seen at *aP2, Ppargc1a*, and *Ucp1* (**Figure 5D-F**). When we examined other genes associated with thermogenesis, such as *Ppargc1a* and *Ucp1* we noted multiple nearby TLE3 binding sites overlapping with putative regulatory regions as detected by DNase hypersensitivity sites (DHS) (**Figures 5E and 5F**). Together, the observations demonstrate that both *Ppargc1a* and *Ucp1* are elevated with the loss of TLE3 (**Figure 3C**), and since these genes are also bound by TLE3 it raises the possibility that TLE3 might be acting to repress their expression.

### TLE3 Represses the Thermogenic Gene Program in iWAT

Thus far we had observed changes in thermogenic genes by real-time PCR *in vitro* and *in vivo* and through ChIP-seq found that TLE3 binds in proximity to these genes, but had only limited insights into how the overall transcriptional landscape of the cells was changing the absence of TLE3. To define the global changes in gene expression provoked by alteration in TLE3 expression, we utilized the tamoxifen-inducible *Tle3*^F/F^ preadipocytes expressing Cre-ERT2. After 4-days of differentiation, we knocked out TLE3 by tamoxifen administration and then assessed changes in gene expression by RNA-sequencing. Consistent with our previous findings, TLE3 deletion led to the up-regulation of genes associated with the thermogenic gene expression program, including *Cidea, Ucp1*, and *Ppargc1a* (**Figure 6A**). Of the 765 genes increased in the TLE3 KO cells, 475 of them may be direct targets of TLE3, as their expression increased in the absence of TLE3 and they have a TLE3 binding site nearby. We note that among these genes there is an overrepresentation of genes that have a role in mitochondrial function (**Figure S6A**). Of the 539 genes which were decreased in the TLE3 KO, 469 of them have nearby TLE3 peaks (**Figure 6B**). These findings are consistent with a dual role of TLE3 in activating or inhibiting gene expression (33).

**Figure 6:**
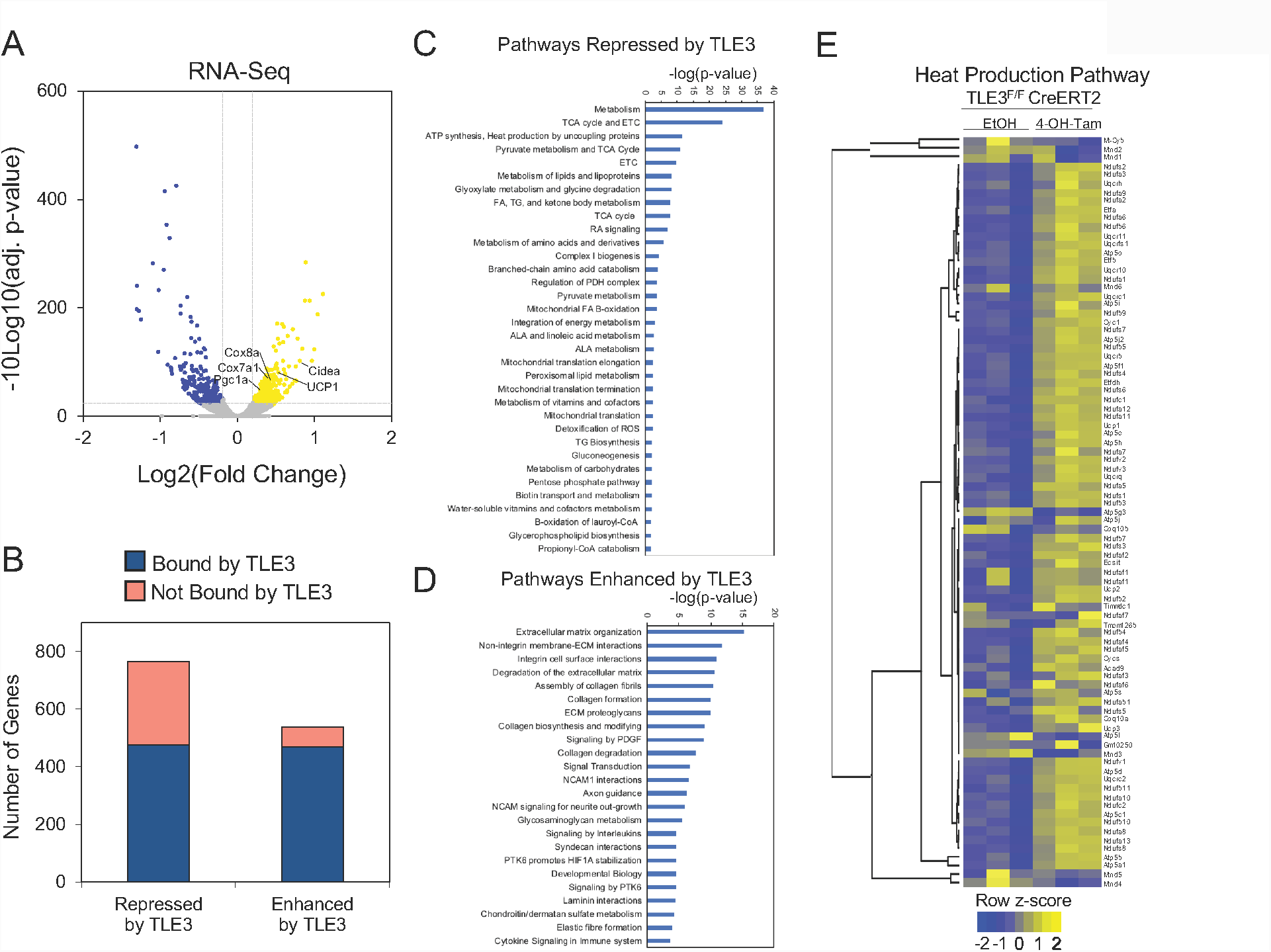
TLE3 represses TCA cycle and thermogenic gene program in iWAT. **(A)** Volcano plot showing differentially expressed genes in preadipocytes isolated from TLE3^F/F^ mice expressing CRE-ERT2. Cells were treated with vehicle or 4-hydroxytamoxifen. (Adjusted p-value < 0.01; Log2 ratio of differential expression > |0.2|) **(B)** Stacked bar graph demonstrating genes that are differentially regulated by TLE3 with or without TLE3 binding site. RNA-seq genes with adjusted p-value < 0.05. **(C)** Gene ontology analysis of 475 genes whose expression is increased in TLE3 KO, and which have at least one TLE3 occupancy site identified by ChIP-seq. Reactome pathways (p<0.0005) listed in order of increasing p-value. **(D)** Gene ontology analysis of 469 genes whose expression is decreased in TLE3 KO, and which have at least one TLE3 occupancy site identified by ChIP-seq. Reactome pathways (p<0.0005) listed in order of increasing p-value. **(E)** Heatmap of genes associated with Reactome pathway R-MMU-163200 (Respiratory Electron Transport, ATP Synthesis by chemiosmotic coupling, and heat production by uncoupling proteins) in TLE3^F/F^ iWAT preadipocytes expressing pMSCV2-CreERT2 and treated with either EtOH (control) or 4-hydroxytamoxifen. Z-score reflects Log2 ratio. (N=3/group)

To characterize the gene program regulated by TLE3, we performed gene ontology analysis using the subset of genes in iWAT that were repressed by TLE3 and had an associated TLE3 site. We found over-representation of genes that encode proteins in metabolic pathways and are involved in energy utilization (**Figure 6C**). Notably, the pathway encompassing heat production by the uncoupling proteins was ranked third highest in significance among those pathways, supporting the notion that TLE3 acts physiologically to repress this program. A heatmap of the changes for all genes included in this pathway is shown (**Figure 6E**). We also performed gene ontology analysis on genes that were enhanced by TLE3 and had an associated TLE3 site. Pathways associated with the extracellular matrix were increased by TLE3, although the statistical significance of this association was lower in comparison to pathways that were downregulated by TLE3 deletion (**Figure 6D**).

### TLE3 Motif Analysis Predicts Functional Interaction with EBFs

Because TLE3 lacks a DNA binding domain, its ability to modulate gene expression depends on interactions with DNA-binding transcription factors. To identify putative transcription factors that recruit TLE3, we completed *de novo* motif analysis on the top 1000 genomic regions identified by TLE3 ChIP-seq, and found sequences that were associated with C/EBP, EBF, FRA1, RUNX, NFIA, TBX20, HOXC9, and ASCL1 motifs (**Figure 7A**). Analysis of individual adipocyte subtypes showed slight variations in the list of transcription factors. Of particular interest was the finding that C/EBP and EBF motifs were consistently highly enriched in sites where TLE3 was bound (**Figure S7A**).

**Figure 7:**
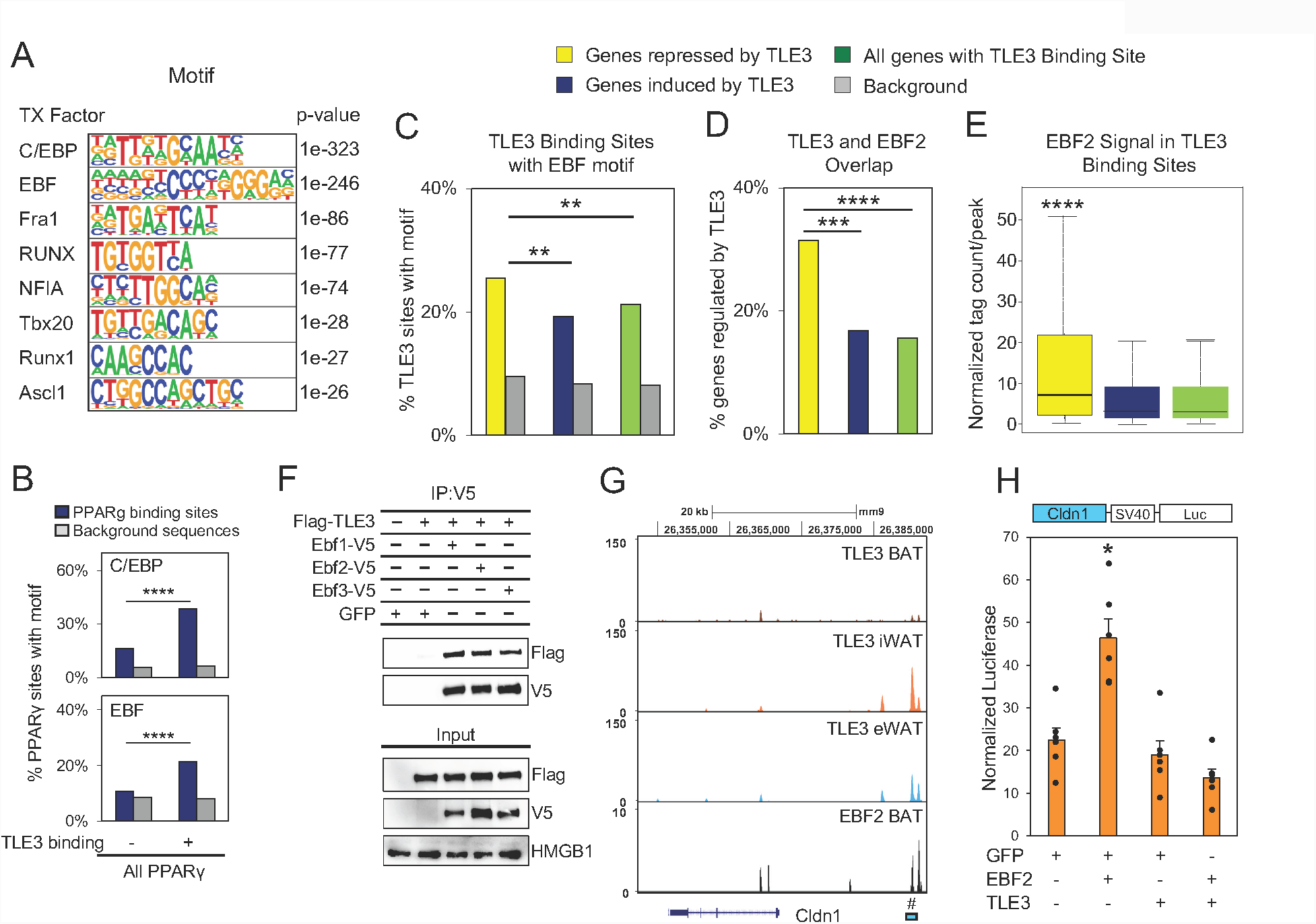
Analysis of TLE3 binding demonstrates repressive interaction with EBF2. **(A)** *De novo* motif analysis of top 1000 TLE3 occupancy sites in WAT, assigned closest matching transcription factor. Listed in order of increasing p-value. **(B)** Known motif analysis of PPARγ binding sites and matching background sequences in the presence or absence of TLE3 WAT binding. **(C)** Known motif analysis of the EBF motif in TLE3 binding sites and matching background sequences within 15kb of the indicated gene groups. **(D)** Overlap between TLE3 and EBF2 binding sites within 15kb of the indicated gene groups. **(E)** EBF2 ChIP-seq signal in TLE3 binding sites within 15kb of the indicated gene groups. **(F)** Coimmunoprecipitation of TLE3 and EBF proteins. 293 cells were transfected with Flag-TLE3 and one of the following: EBF1-V5, EBF2-V5, or EBF3-V5. Lysates were immunoprecipitated with anti-V5 antibody and the precipitates were analyzed by immunoblotting. **(G)** Genome browser tracks of Cldn1 showing overlapping peaks between TLE3 and Ebf2. # indicates genomic region cloned into luciferase reporter assay. **(H)** Analysis of Cldn1 enhancer activation of luciferase reporter by coexpression of EBF2, TLE3, or both in beige preadipocytes differentiated *in vitro*. Data presented as mean ± SEM. Significance analyzed by two-tailed Student’s t-test (Figure 7H), Fisher’s exact test (Figures 7C and 7D), or Wilcoxon rank sum test (Figure 7E). *p<0.05, **p<0.01, ***p<2.2e-8, ****p<2.2e-16.

To understand the mechanisms by which TLE3 modulates the recruitment of beige adipocytes in inguinal white adipocytes, we analyzed TLE3-binding sites that overlap with PPARγ. We found that the C/EBP and the EBF motif were enriched at PPARγ sites that are associated with TLE3 (**Figure 7B**). Given the potential cross-talk of these transcription factors, we next analyzed putative target genes of TLE3 in iWAT adipocytes (i.e. genes that were repressed or induced by TLE3 KO and display nearby TLE3 binding) (**Figure S7B**). Interestingly, the EBF motif was present in a higher fraction of TLE3 binding sites near genes that were repressed by TLE3 compared to genes that were enhanced by TLE3 (**Figure 7C**). This was not observed for the C/EBP motif, where we found a similar number of C/EBP motifs in genes stimulated or repressed by TLE3 **(Figure S7C).** Interestingly, some of the transcripts regulated by TLE3 have been reported to be direct targets of EBF2 (28). We therefore analyzed the coincidence of TLE3 and EBF2 binding near genes regulated by TLE3 on a genome-wide level. We compared a ChIP-seq datasets from EBF2 in BAT (30) with the TLE3-binding profile in iWAT and found a greater overlap between TLE3 and EBF2 binding sites near genes that were repressed by TLE3 compared to genes enhanced by TLE3 (**Figure 7D**). The binding intensity of EBF2 was higher in TLE3-binding sites near repressed genes (**Figure 7E**), suggesting that crosstalk between TLE3 and EBF2 primarily occurs near genes directly repressed by TLE3.

To test the functional connection between TLE3 and EBFs, we explored interactions between TLE3 and EBF1, EBF2 and EBF3. Cells were transfected with Flag-TLE3 in the presence or absence of V5-tagged EBF proteins, immunoprecipitated with anti-V5 antibody, and immunoblotted with anti-Flag antibody. We found that TLE3 co-immunoprecipitated with EBF1, EBF2, and EBF3 (**Figure 7F**). Because we observed that TLE3 and EBF2 colocalized near several putative target genes, we sought to investigate how TLE3 influenced the transcriptional activity of EBF2. We utilized the ChIP-seq data of TLE3 and EBF2 (39) to identify a site where both TLE3 and EBF2 colocalized in proximity to the BAT-selective marker, CLDN1 (**Figure 7G**). 293T cells were transfected with a luciferase reporter driven by the 0.5 kb enhancer element upstream of *Cldn1* gene. This enhancer element was defined as a DHS site that has overlapping TLE3 and EBF2 occupancy (**Figure S7D**). Beige adipocytes were also transfected with an EBF2 expression vector, TLE3 expression vector, or both. We found that EBF2 activated the *Cldn1* reporter and that reporter expression was inhibited to baseline levels when TLE3 was also expressed (**Figure 7H**). These findings demonstrate that TLE3 can counter EBF2 transcriptional activity.

## Discussion

Subcutaneous white adipose tissue has a tremendous capacity to change its morphology in response to the cold. Mice that are acclimated to the cold acquire thermogenic cells that are rich in mitochondria and are believed to play a long-term adaptive role in thermogenesis. The mechanisms that regulate the mitochondrial gene expression program in beige adipocytes are incompletely understood. Here we have outlined a role for the transcriptional cofactor TLE3 in regulating the mitochondrial beige adipocyte program. The loss of TLE3 in adipocytes promotes the recruitment of beige adipocytes in the inguinal white adipose tissue depot in response to long-term cold exposure, as shown by gross histological analysis, transcriptional changes, and increased expression of UCP1. Our findings further support the concept that increasing beige adipocyte number can lead to improvements in glucose metabolism and insulin sensitivity. We demonstrated that TLE3 regulates the mitochondrial gene expression program in beige adipocytes and provide evidence indicating that this occurs through countering EBF2 action, a transcription factor that promotes the recruitment of thermogenic cells (28, 40). These studies highlight that mechanisms that promote the recruitment of beige adipocytes can prevent weight gain associated with aging.

Our findings strongly support the contention that promoting an increase in beige adipocyte activity is sufficient to have beneficial effects on systemic glucose metabolism. We found that loss of TLE3 improved glucose metabolism in the setting of cold exposure, but not at thermoneutrality, and that this effect was correlated with the abundance of beige adipocytes. We found differences in histology and gene expression consistent with beige adipocytes between control and TLE3-deficient mice during cold exposure but not thermoneutrality, strongly suggesting that increasing the amount of beige fat alone can lead to improvements in glucose metabolism. When testing to see which tissues were primarily involved in clearing glucose, we found that the inguinal white adipose tissue depots had the greatest increase in glucose uptake. Thus, despite the fact that they represent a small fraction of the total adipocyte population, beige adipocytes can exert marked effects on systemic energy balance.

Our data suggest that the increase in glucose utilization provoked by TLE3 deletion is likely due to increased expression of glucose transporters and genes involved in mitochondrial oxidative phosphorylation. Gene ontology analysis pointed to broad changes in expression in metabolic pathways in the absence of TLE3, including components of the ETC, heat production, fatty acid oxidation, and the TCA cycle. Loss of TLE3 also led to enhanced energy expenditure under conditions where mice had more beige adipocytes. In the cold, mice lacking TLE3 in adipocytes lost more weight, an outcome that was largely attributed to increased energy expenditure. This was also seen *in vitro*, where beige adipocytes lacking TLE3 had increased basal respiration and uncoupling. When mice were challenged with a HFD, they were protected against diet-induced weight gain even at ambient temperature.

In both rodents and humans, aging can lead to an increase in adiposity, and impaired glucose metabolism (41, 42). However, there is little known about the mechanisms that drive this age-induced weight gain. We suspect that the loss of beige adipocytes that occurs with aging (16), likely contributes to this weight gain. We found that loss of TLE3 blunted the accumulation of adipose tissue mass in older mice. While adipose mass doubled when control mice were aged in our study, the fat mass in young and aged TLE3-deficient mice was similar. These findings identify TLE3 as a contributor to the age-dependent weight gain seen in rodents.

Although TLE3 has been studied broadly during development, there has heretofore been little known about its genome-wide distribution in adipocytes. By integrating RNA-seq and ChIP-seq analysis we identified putative genomic targets of TLE3. Most of these were found in distal genomic regions that were greater than 25kb from the TSS. Given that TLE3 lacks a DNA-binding domain, we performed motif analysis on genomic regions associated with TLE3 and found that C/EBP and EBF motifs were highly associated with TLE3 binding. Our analysis also supports an overlap in genome-wide occupancy of TLE3 and PPARγ, where 25% of TLE3 binding sites were associated with PPARγ, while 49% of PPARγ binding sites overlap with TLE3. After analyzing TLE3-binding sites near genes regulated by TLE3 in beige adipocytes, we found that EBF2 and TLE3 colocalized near genes activated with the loss of TLE3. This led us to investigate the crosstalk between TLE3 and EBFs. We found that TLE3 interacted with EBF2, repressed EBF2 target genes, and repressed EBF2 transcriptional activity in a reporter assay. Further, we found that genes that are repressed by TLE3 were more likely to contain an EBF motif compared to those that were enhanced by TLE3. There was a high degree of overlap between TLE3- and EBF2-binding sites near genes that were repressed by TLE3, and the TLE3 binding intensity was consistently higher at sites near TLE3-repressed genes. These findings indicate a new mechanism by which TLE3 attenuates the beige adipocyte gene expression program by serving as a molecular brake on EBF2.

In conclusion, we have identified TLE3 as a negative regulator of EBF2 and the beige adipocyte program, whose actions block energy expenditure, glucose utilization, and mitochondrial energetics. Developing therapeutics that target TLE3 may provide opportunities to prevent weight gain associated with diet and aging.

## METHODS

### Animals

A conditional knockout allele for *Tle3* was generated on a 129/OlaHsd and C57BL/6J background as has been described (34). TLE3^flox/flox^ mice were bred for 9 generations to C57BL/6J background. To obtain age-matched littermates, TLE3^flox/flox^ mice were bred to TLE3^flox/flox^ Adipoq-Cre BAC transgenic mice bred for 9 generations by the Jackson laboratories to C57BL/6J background. Except where otherwise stated, male mice, age 3-4 months, that expressed the Adipoq-Cre transgene were compared to male littermate controls lacking the Cre transgene.

C57BL/6J mice for aging studies were obtained from the National Institute of Aging, courtesy of the Donato Lab.

Mice were housed at 22-24°C under a 12-hour light/dark cycle with *ad libitum* access to food and water, except when food was withdrawn for an experiment. Animals were maintained on a Teklad global soy protein-free diet (2920x), except where otherwise noted.

### Cell Culture

Preadipocytes isolated from the stromal vascular fraction of inguinal white fat depots were isolated as previously described (43); cells were immortalized by retroviral expression of the SV40 large T antigen (Cat#13970, Addgene) using a hygromycin selection marker, and expressing MSCV CreERT2 (Cat#22776, Addgene) using a puromycin selection marker. Adenovirus was amplified using Phoenix-ECO cells. Adenovirus vectors were purified by CsCl gradient.

Preadipocytes were plated in a 12-well plate in DMEM, 10% FBS, 20 nM insulin, 1nM T3, with a seeding density of 0.1 x 10^6^ cells per well. Upon confluence, cells were differentiated using DMEM media containing 10% Fetal Bovine Serum (FBS; Cat#FBS-BBT-5XM, RBMI), 20nM insulin, 1nM T3, 0.5 mM isobutylmethylxanthine, 0.5 μM dexamethasone, 0.125 mM indomethacin and 1 μM rosiglitazone. After two days the media was changed to DMEM, 10% FBS, 20 nM insulin, 1nM T3 and 1 μM rosiglitazone. On the fourth day cells were either given 500 nM Tamoxifen to induce knockout of TLE3, or an equivalent volume of ethanol, as a control.

### Cell Staining

After 10 days of differentiation, cells were washed three times in 1x PBS and fixed with freshly prepared PBS + 4% formaldehyde for 1 hour, then incubated with DAPI (D3571, Thermo Fisher) and 1 μg/ml BODIPY (D3922, Thermo Fisher) for 20 minutes. Cells were subsequently washed three times in 1x PBS and visualized by GFP and DAPI channels.

Following DAPI and BODIPY visualization, cells were stained using stock solution Oil Red O (0.5g in 100 mL isopropanol; O0625, Sigma Aldrich) diluted to a freshly-prepared working solution (6 mL stock solution and 4 mL ddH_2_O) solution, and incubated for 1 hour.

### Western Blot Analysis

Cells were homogenized using Radioimmunoprecipitation assay (RIPA) buffer (Boston Bioproducts, Inc.) plus proteinase inhibitor cocktail (Cat# 04693124001, Sigma Aldrich) and lysed using a 25-gauge needle. Protein was extracted from flash-frozen tissues in RIPA buffer with protease inhibitor cocktail using a glass dounce. Cell or tissue homogenate was spun at 10,000 x *g* for 10 min at 4°C, and the concentration of the supernatant was measured using the Pierce BCA Protein Assay Kit (Cat# 23225, Thermo Fisher). Protein samples (10-20ug per lane) were denatured in Laemmli loading buffer, heated at 70°C for 20 min, resolved by SDS-PAGE gels (typically 10%), and transferred to nitrocellulose membranes (GE Healthcare). Membranes were probed for TLE3 (Cat#11372-1-AP, Proteintech; Cat#ab213596, Abcam), UCP1 (Cat#ab10983, Abcam), PPARγ (Cat#22061-1-AP, Proteintech), β-actin (Cat#4970, Cell Signaling; Cat#3700, Cell Signaling), HMGB1 (Cat#ab18256, Abcam), V5 (Cat#46-0705, Thermo Fisher), or Flag M2 (Cat#F3165, Millipore-Sigma).

For analysis of mitochondrial oxidative phosphorylation proteins, protein samples were denatured in Laemmli loading buffer, heated at 50°C for 20 min, resolved by SDS-PAGE gels (5-20% gradient, BioRad), and transferred to a PVDF membrane (BioRad) in CAPS buffer (10mM CAPS, 10% methanol, pH 11.0). Membrane was probed using the Total OXPHOS Rodent Antibody Cocktail (Cat#ab110413, Abcam).

### Gene Expression

Total RNA was isolated using TRIZOL reagents (Cat# 15596018, Thermo Fisher) and reverse transcribed using SS VILO Mastermix (Cat# 11755500, Thermo Fisher). Gene expression of TLE3 was quantified using TAQMAN Universal Master Mix II (Cat# 4440038, Applied Biosystems) and Quant Studio 6 instrument (Applied Biosystems by Invitrogen). All other transcripts were quantified by real-time qPCR using KAPA SYBR FAST Rox Low (Cat# KK4621, Kapa Biosystems). Primer sequences are shown in Table S1.

### In Vitro Glucose Uptake

TLE3^F/F^ iWAT preadipocytes were plated in a 12-well plate, allowed to reach confluence, then stimulated to differentiate. After two days, the media was changed to DMEM, 10% FBS, 20 nM insulin, 1 nM T3 and 1 μM rosiglitazone. On the fourth day cells were either given 500 nM Tamoxifen to induce knockout of TLE3, or an equivalent volume of ethanol, as a control. On the tenth day, the cells were incubated in DMEM, 10% FBS, 20 nM insulin, and 1 nM T3 (no rosiglitazone) for 3 hours at 37°C. Cells were washed twice in warm 1x PBS solution, and incubated with 0.1 uCi of 2-[^14^C(U)]-Deoxy-D-glucose (Cat# NEC720A050UC, Perkin Elmer) per well, with unlabeled 2-deoxy-D-glucose up to 6.5 mM total glucose concentration. Negative control wells were also given 20 uM cytochalasin B. After a five-minute incubation cells were washed 3 times in 1x PBS solution at 4°C, then 1 mL PBS with 1% SDS was added to each well to lyse the cells. After 10 minutes incubation, cell mixture was mixed by pipette. A sample was taken for measuring protein concentration, and another sample was taken to measure radioactivity by liquid scintillation counter in Ultima Gold MV (Cat#6013159, Perkin Elmer). Scintillation counts were normalized to total protein concentration.

### Dual-Luciferase Reporter Assay

iWAT preadipocytes were plated and differentiated with the differentiation cocktail described above. Four days after differentiation, the cells were trypsinized and plated in a 48-well plate with a seeding density of 20,000 cells/well. After 24 hours, the cells were co-transfected with 50 ng pGL3-firefly luciferase reporter vector (Cat# E1960, Promega) driven by the Cldn1 enhancer element, 1 ng *Renilla* luciferase vector (Cat# E1960, Promega) and separate expression vectors containing 50 ng GFP, 50 ng Ebf2, and/or 100 ng TLE3. Cells were transfected using Lipofectamine 3000. The next day cells were washed and lysed. 20uL of cell lysate was incubated with the luciferase assay substrate and both firefly and *Renilla* luciferase activities were measured. *Renilla* luciferase activity was used as an assay control.

### Prolonged Cold Exposure and Thermoneutrality

For cold experiments, male mice, age 3-4 months were put at 2-8°C for 4 days. For thermoneutrality experiments, male mice, age 3-4 months were put at 30°C for one week.

### Energy Expenditure

Energy expenditure, foot and water intake, and ambulatory activity were determined by using Comprehensive Lab Animal Monitoring System (CLAMS) (Columbus Instruments). Animals were housed individually within the metabolic cages on a 12-hour light/dark cycle with ad libitum access to food (chow diet) and water. System temperature was set at 4°C and remained at 4-5°C throughout the run. Energy expenditure was calculated as a function of oxygen consumption and carbon dioxide production in the CLAMS cages. Energy expenditure = (3.815 + 1.232*(VCO_2_/VO_2_))*VO_2_.

### Histology

Adipose tissue was fixed in 4% buffered formaldehyde was fixed in 4% buffered formaldehyde and embedded in paraffin. Sections were stained with hematoxylin and eosin using standard protocols.

### Glucose and Insulin Tolerance Tests

A glucose tolerance test (GTT) was administered by fasting mice for 6 hours with free access to water during fasting, and fasting blood glucose levels was measured using a glucometer by tail bleeds. Mice were then given an intraperitoneal injection of glucose (1mg/g). Blood glucose levels were measured in intervals (0, 15, 30, 60, 120 min post injection) to determine how quickly the glucose is cleared from the blood.

An insulin tolerance test (ITT) was administered in a similar manner, but using intraperitoneally injected insulin (Cat# NDC 0002-8215-17 HI-213, Lilly) at 0.75U/kg.

### CL-316,243 Treatment

TLE3^F/F^ Adipoq-Cre male mice and their littermate controls were housed at 30°C to control for endogenous BAT activation. Mice were injected intraperitoneally with 1 mg/kg/day of CL-316,243 (CAS 138908-40-4, Cayman Chemical) for 3 days, or equivalent volume of 1x PBS. Prior to and after 3 days of CL-316,243 administration, animals were fasting for 6 hours and fasting blood glucose was measured via tail vein.

### In Vivo Glucose Uptake

Male TLE3^F/F^ Adipoq-Cre mice and their littermate controls were placed at 2-8°C for 4 days with free access to food and water. On the fourth they, mice were fasted for 6 hours and given a tail-vein injection of 4 uCi of 2-[^14^C(U)]-dexoxy-D-glucose (Cat# NEC720A050UC, Perkin Elmer) in 1x PBS. Blood samples were taken at 0, 5, 10, 20, and 30 minutes to monitor clearance of radioactive glucose from the blood. After 30 minutes, animals were sacrificed, tissues were weighed and dissolved in 1 mL of NaOH at 60°C for one hour. Samples were neutralized. One part of homogenate was used to determine content of unphosphorylated glucose. This sample was deproteinated by adding equivalent volumes of 0.3 M Ba(OH)_2_ and 0.3 M ZnSO_4_, centrifuged, and the ^14^C content of the supernatant was measured by liquid scintillation counting. Another part of the homogenate was used to determine content of total glucose. Insoluble proteins in this sample were removed by adding 6% HClO_4_, centrifuged, and the ^14^C content of the supernatant was measured by liquid scintillation counting. The amount of phosphorylated 2-[^14^C(U)]-Deoxy-D-glucose was then determined by subtracting the radioactivity counts of the unphosphorylated sample from radioactivity counts of the total sample and normalized to tissue weight.

### High Fat Diet

Where noted, male Adipo-TLE3 KO and littermate controls were put on 60% high fat diet (HFD) (D12492, Research Diets) starting at 2 months of age, and maintained for 14 weeks. Weight was measured weekly.

### Body Composition/MRI Measurements

Composition of lean tissue, fat mass, and fluid was analyzed using the Bruker minispec LF50 Body Composition Analyzer.

### TLE3 ChIP-seq

Cells from the stromal vascular fraction (SVF) of brown adipose tissue (pooled interscapular, cervical, and axillary depots) as well as the epididymal (eWAT) and inguinal (iWAT) adipose tissue of male NMRI mice were obtained and differentiated with rosiglitazone as previously described (23). Mature adipocytes were analyzed at day 7, and only cultures with at least 50% degree of differentiation were used.

For preparation of material for TLE3 ChIP-seq, eWAT-, iWAT-, and BAT-derived adipocytes were treated with 2 mM disuccinimidyl glutarate (DSG) (Proteochem) for 15 min followed by cross-linking with 1% formaldehyde for 30 min. Crosslinking was stopped by addition of glycine. Nuclei isolation, chromatin preparation and chromatin IP was performed as described previously (23). The chromatin IP was performed using anti-TLE3 (Proteintech 11372-1-AP). ChIP-seq libraries were constructed according to the manufacturer’s instructions (Illumina) as described in (44).

### TLE3 ChIP-seq data analysis

ChIP-seq data sets were mapped to the mm9 genome with STAR (45) set to not map across splice junctions, as previously described (46), and tag directory generation, peak calling/annotations and motif analyses were performed with HOMER (47). TLE3 binding sites were called using “-style factor” and “-F 10” settings with all other parameters set at default in eWAT-derived, iWAT-derived and BAT-derived adipocytes, and overlapping peaks were merged into one master peak file. Subsequently, TLE3 peaks in each condition were identified as having >15 normalized tags/peak in a 230bp window and 10-fold more tags relative to the input control. For known and *de novo* motif analyses we used HOMER’s findMotifsGenome.pl with default settings in the indicates groups of TLE3 binding sites. For comparison with PPARγ and EBF2 binding profiles, datasets were obtained from the Gene Expression Omnibus (GEO) under accession numbers GSE41481 and GSE97116, respectively. Mapping and peak calling was performed with a similar strategy as for TLE3 ChIP-seq data. PPARγ profiles were divided into binding sites (>25tags/peak and 10-fold more tags than input) with high TLE3 binding (i.e. 25 tags/window and 10-fold more tags than input) and no TLE3 binding (<15 tags/window). To determine genomic cooccurence of EBF2 and TLE3 binding, we defined TLE3 binding sites near the indicated gene groups and used HOMER’s mergePeaks program to count the literal overlaps of binding sites. Furthermore, EBF2 tag counts were summarized in a 230bp window in TLE3 binding sites near indicated gene groups.

The University of California at Santa Cruz (UCSC) Genome Browser (48) was used for visualization of genome tracks.

### EBF2 ChIP-seq for Genome Tracks

EBF2 ChIP-seq data from immortalized BAT preadipocytes differentiated two days in cultures were obtained from GSM1913014 (39).

### Gene Expression Profile - Input Data

Intact poly(A) RNA was purified from total RNA samples (100-500 ng) with oligo(dT) magnetic beads and stranded mRNA sequencing libraries were prepared as described using the Illumina TruSeq Stranded mRNA Library Preparation Kit (RS-122-2101, RS-122-2102). Purified libraries were qualified on an Agilent Technologies 2200 TapeStation using a D1000 ScreenTape assay (cat# 5067-5582 and 5067-5583). The molarity of adapter-modified molecules was defined by quantitative PCR using the Kapa Biosystems Kapa Library Quant Kit (cat#KK4824). Individual libraries were normalized to 10 nM and equal volumes were pooled in preparation for Illumina sequence analysis.

Sequencing libraries (25 pM) were chemically denatured and applied to an Illumina HiSeq v4 single read flow cell using an Illumina cBot. Hybridized molecules were clonally amplified and annealed to sequencing primers with reagents from an Illumina HiSeq SR Cluster Kit v4-cBot (GD-401-4001). Following transfer of the flowcell to an Illumina HiSeq 2500 instrument (HCSv2.2.38 and RTA v1.18.61), a 50 cycle single-read sequence run was performed using HiSeq SBS Kit v4 sequencing reagents (FC-401-4002).

### Gene Expression Profile – Analysis

Fastq sequences were aligned with Novoalign (v. 2.8) against the mouse genome (UCSC release mm9) with splice junctions. Alignments to splice junctions were converted to genomic coordinates with USeq SamTranscriptomeParser (version 8.8.8). RNASeq quality was evaluated by running Picard CollectRnaSeqMetrics. Differential gene expression was identified using the USeq application DefinedRegionDifferentialSeq (version 8.8.8), which counts alignment tags over collapsed Ensembl gene annotation, executes the R DESeq2 application, filters the gene results at log2 ratio of 1.0 and −10log10(adjusted P) of 20, and reports the results.

The resulting gene lists were used for functional enrichment analysis (geneontology) using the Panther Classification System (49) and Reactome pathway databases (50).

### Data Availability

RNA-sequencing data are available from the Gene Expression Onmibus (GEO) under accession number GSE116894. ChIP-sequencing data are available from the Gene Expression Omnibus (GEO) under accession number GSE116767.

### Statistical Analysis

Data are presented as mean ± SEM, unless otherwise stated. Student’s t test was used to determined significance unless otherwise stated.

### Study Approval

Animal experiments were conducted with the approval of the Institutional Animal Care and Use Committee (IACUC) of the University of Utah.

## AUTHOR CONTRIBUTIONS

S.P., A.L., P.R., J.S., S.L. and C.J.M. performed experiments. S.P., A.L, P.R., P.T., S.M., and C.J.V conceived of experiments and analyzed data. S.P. and C.J.V wrote manuscript. P.T., S.M., A.D., and C.J.V. provided resources. C.J.V. secured funding.

## ACKNOWLEDGEMENTS

We would like to thank members of the Diabetes and Metabolism Center and the Biochemistry Department at the University of Utah for useful discussion and feedback. We are grateful to the Metabolic Phenotyping Core, a part of the Health Sciences Cores at the University of Utah for work performed and access to instrumentation, as well as the High-Throughput Genomics Core, at the Huntsman Cancer Institute of the University of Utah for their assistance with performing RNA-sequencing and data analysis. This study was supported by 1R01DK103930, R03DK103089, K01DK097285, DRC, P30DK020579, 5T32DK091317, and P30CA042014. Work in the Mandrup laboratory was supported by grants from the Lundbeck Foundation, the Novo Nordisk Foundation, and the Danish Independent Research Council | Natural Sciences. The content is solely the responsibility of the authors and does not necessarily represent the official views of the National Institute of Diabetes and Digestive and Kidney Diseases (NIDDK), National Cancer Institute (NCI) or the National Institutes of Health (NIH).

